# BERTax: taxonomic classification of DNA sequences with Deep Neural Networks

**DOI:** 10.1101/2021.07.09.451778

**Authors:** Florian Mock, Fleming Kretschmer, Anton Kriese, Sebastian Böcker, Manja Marz

## Abstract

Taxonomic classification, i.e., the identification and assignment to groups of biological organisms with the same origin and characteristics, is a common task in genetics. Nowadays, taxonomic classification is mainly based on genome similarity search to large genome databases. In this process, the classification quality depends heavily on the database since representative relatives have to be known already. Many genomic sequences cannot be classified at all or only with a high misclassification rate.

Here we present `BERTax`, a program that uses a deep neural network to precisely classify the superkingdom, phylum, and genus of DNA sequences taxonomically without the need for a known representative relative from a database. For this, `BERTax` uses the natural language processing model BERT trained to represent DNA. We show `BERTax` to be at least on par with the state-of-the-art approaches when taxonomically similar species are part of the training data. In case of an entirely novel organism, however, `BERTax` clearly outperforms any existing approach. Finally, we show that `BERTax` can also be combined with database approaches to further increase the prediction quality.

Since `BERTax` is not based on homologous entries in databases, it allows precise taxonomic classification of a broader range of genomic sequences. This leads to a higher number of correctly classified sequences and thus increases the overall information gain.

## 1 Introduction

How do we know what kind of organisms we have sequenced?

This question seems at first rather strange, as traditionally, DNA sequencing was mostly performed on cultivated cells or viral strains/isolates. However, in recent years, this has become a common problem, especially due to metagenomics, where genetic material is directly recovered from environmental samples and includes unknown compositions of organisms. To answer this question we need to classify the taxonomic origin of the sequences.

For this task of taxonomic classification, it became common to use a homology based approach of DNA/RNA sequences queried against databases. Such approaches achieve a high level of precision and can even determine the exact species when the genome is already known and present in the database.

However, many query sequences cannot be classified at all, and therefore only a low recall is achieved. One of the reasons is, that only a fraction of all species (at most 14 % of all eukaryotic terrestrial and 9 % of all ocean species) have been described and only a fraction of these are stored as reference genomes in current databases [1, 2]. Furthermore this estimate ignores bacteria and viruses for which the situation is likely way worse [2].

On the technical level, most taxonomic classification tools use either (a) local alignments, for example MegaBLAST [3] and DIAMOND [4], (b) *k*-mers as in CLARK [5] and CDKAM 2 [6], (c) are based on Burrow-Wheeler transformations, either on DNA level (`centrifuge` [7]) or on protein level (`Kaiju` [8]). Also commonly used are hybrid methods, such as a combination of (d) local alignments and *k*-mers such as k-SLAM [9] and `MMseqs2` [10] (e) or a combination with minimizers: based on *k*-mers like `Kraken2` [11] and `sourmash` [12] or based on local alignments like `minimap2` [13].

Using local alignments is a very precise but relatively slow method. These methods typically need a seed region with high similarity, typically resulting in a limited recall, meaning that only a portion of the samples can be predicted. The usage of *k*-mers or minimizer requires even wider regions with high similarity, which can result in an even lower recall and precision. However *k*-mers and minimizer are significantly faster [14]. Overall, the taxonomic classification quality depends mainly on the quality of the database and less on the methodology of the database usage [14].

Recently multiple deep neural network approaches were developed to overcome some of the described limitations.

Instead of having to rely on similar sequences being present in a database, deep learning methods allow modeling complex dependencies between the data and the target variable, in our case, the DNA sequences and corresponding taxonomic class. Typically for deep learning methods, however, interpretability is often not an easy task.

One example for a deep learning approach is `DeepMicrobes` [15], which is specifically designed for taxonomic classification of microbial data from human gut, however, it can be used as generic classifier for any DNA classification task. It is important to mention that `DeepMicrobes` relies on *k*-mer embeddings (k=12), similar to non-machine learning methods in handling the sequence data.

Here, we present the tool `BERTax` for classification of DNA sequences on three different taxonomic levels, superkingdom (archaea, bacteria, eukaryota and viruses), phylum, and genus. The fundamental novelty is to assume DNA as a “language” and to classify the taxonomic origin based on this language understanding rather than by local similarity to known genomes in a database.

As a result we achieve a classifier not limited to typical restrictions as comparable tools, as `BERTax` is not limited to coding regions, or specific superkingdoms like bacteria or viruses, but can potentially classify any genome region, from any of the four superkingdoms and does not require similar sequences in the database.

As basis to our model, we use the architecture BERT (Bidirectional Encoder Representations from Transformers) [16], which is a state-of-the-art model for many natural language processing tasks [17]. Being standard for BERT, we (a) pre-trained the deep neural network (DNN) and (b) fine-tuned the DNN. In this approach, to classify a query sequence, less exact representatives are needed in the training data since the training sequences are not memorized *per se* [16].

We developed three different DNN architectures: the (1) flat, (2) nested, and (3) all-in-one architecture, which we compared with each other. The latter and best performing architecture has been compared with common database approaches and the deep-learning approach `DeepMicrobes`. The performance of `BERTax` can further be increased by combining a precise database approach, like `MMseqs2` with `BERTax`.

The final DNN model used in `BERTax` outperforms any other tool when unknown genera are used as input data: average precision (AveP) can be increased from 67.69 % to 87.94 % on the level of superkingdoms and from 43.62 % to 59.06 % on phylum-level. But also when closely related organisms are used for training, `BERTax` classifies genomic sequences on average with a slightly higher precision than previous tools.

## 2 Methods

`BERTax` is based on BERT [16], a state-of-the-art model for many natural language processing (NLP) tasks. BERT is a deep neural network (DNN) encoder architecture that relies on a transformer employing the mechanism of self-attention [18]. Self-attention is a method determining autonomously which parts of the inputs are relevant to each other. This enables the transformer architecture to process the sequential data not in a predefined order, enabling a faster training process and therefore the recognition of even more complex interrelationships in the same time [18].

BERT appears to outperform previous methods due to the division of the training process into unsupervised pre-training and supervised fine-tuning, and due to its deeply bidirectional nature of language processing [16]. Deeply bidirectional means that both reading directions are considered simultaneously, as opposed to the “shallowly” bidirectional approach where the model is trained on both directions independently, as used in the competing method ELMO (Embeddings from Language Models) [19, 20].

### 2.1 Pre-training: learning the “DNA language”

During pre-training, the model is trained on two tasks that are not specific to the actual classification objective, namely Masked Language Modeling (MLM) and Next Sentence Prediction (NSP). MLM masks individual tokens (in NLP a token is a word or a part of a word) which are then to be predicted. NSP predicts whether the second of two sentences is related to the first or whether it was replaced by a random independent sentence. When training the BERT model, MLM and NSP are trained together, with the goal of minimizing the combined loss function of the two strategies [16].

To model DNA as natural language, `BERTax` was pre-trained on DNA sequences of a fixed length of 1, 500 nucleotides. Each sequence was split into 500 3-mers (i.e., tokens in a natural language), with the first 250 3-mers being the first sentence and the second 250 3-mers being the second sentence (see Fig. 1). For each input, 15 % of the 3-mers were masked for MLM to learn what token fits which context and for 50 % of the inputs, the second sentence was randomly replaced by a sentence from another DNA fragment as a negative training set for NSP only.

**Figure 1:**
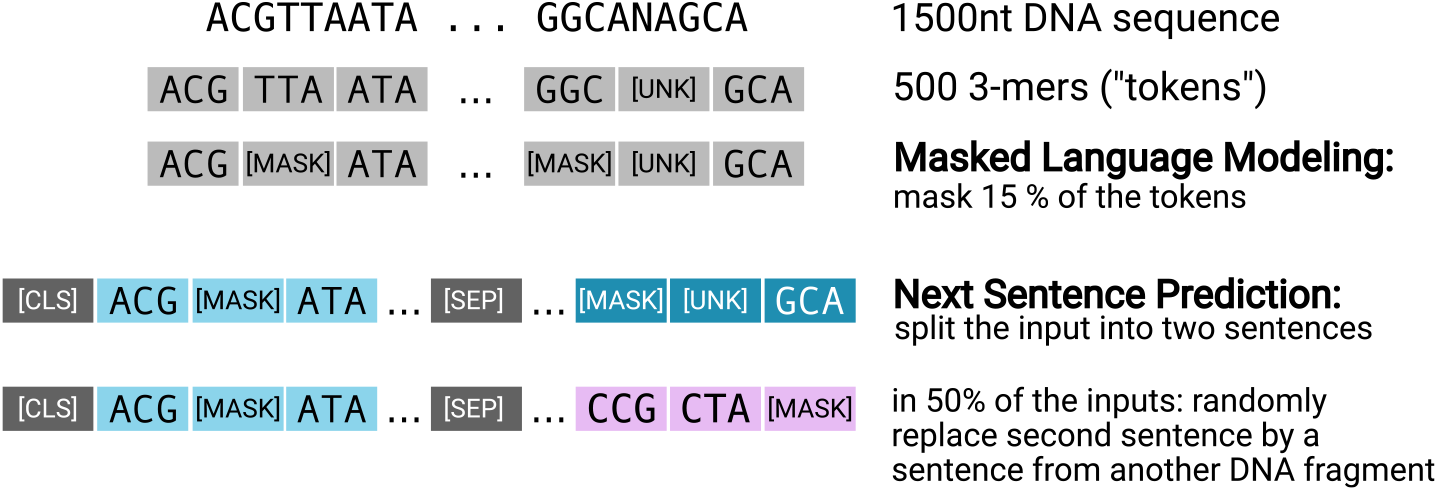
Pre-training of BERT on DNA. DNA sequences of length 1500 are divided into 500 non-overlapping tokens (each representing a 3-mer), of which 15 % are randomly masked so that the neural network can no longer know the original token. Further, the 500 tokens are divided into two sentences of length 250 tokens, of which the second sentence is randomly replaced 50 % of the time by a sentence from a different sequence. The task of predicting whether the sentence has been replaced or not is called Next Sentence Prediciton (NSP), the task of predicting the masked tokens is called Masked Language Model (MLM). Internally, in our BERT model, each token is represented by a vector of length 250.

Internally, a 3-mer is a token (i.e., a string with an assigned meaning), with 64 possible tokens assuming the occurrence of the four canonical nucleotides. BERT uses additionally five specific tokens necessary for training: the unknown token `[UNK]`, representing all 3-mers containing at least one ambiguously sequenced character, such as “`N`”; the empty token `[PAD]` which is required for padding shorter input sequences to the required input length of 500 tokens; the classification token `[CLS]`, which is designed to represent the “meaning” of the entire sentence; the mask token `[MASK]`, which masks the words to be predicted in MLM; and the separator token `[SEP]`, which separates the two sentences for the NSP task.

On the highest hierarchical level, the BERT architecture consists of a specified number of layers called transformer blocks (see Fig. 2A). Each transformer block contains a certain number of attention heads – where weights are learned using the mechanism of self attention – and one feed-forward layer, which serves as the connection between the transformer blocks. Hyper-parameters, i.e., parameters whose values are used to control the learning process, include the number of transformer blocks (set to 12), the number of attention heads per block (5) and the size of the feed-forward layers (1024). The embedding dimension, i.e., the dimension of the internal representation of the sequences, is set to 250 (see Fig. 1). The dropout rate (fraction of nodes not trained per epoch) of the feed-forward layers, used for better generalization, is set to 5 %. All hyper-parameters are set to lower values than those used in the original BERT models [16]. The main reason for this is, that the vocabulary size for the 3-mer DNA sequences is much lower – 65 as opposed to approximately 30,000 for the English language tokens used in the original BERT models [21].

**Figure 2:**
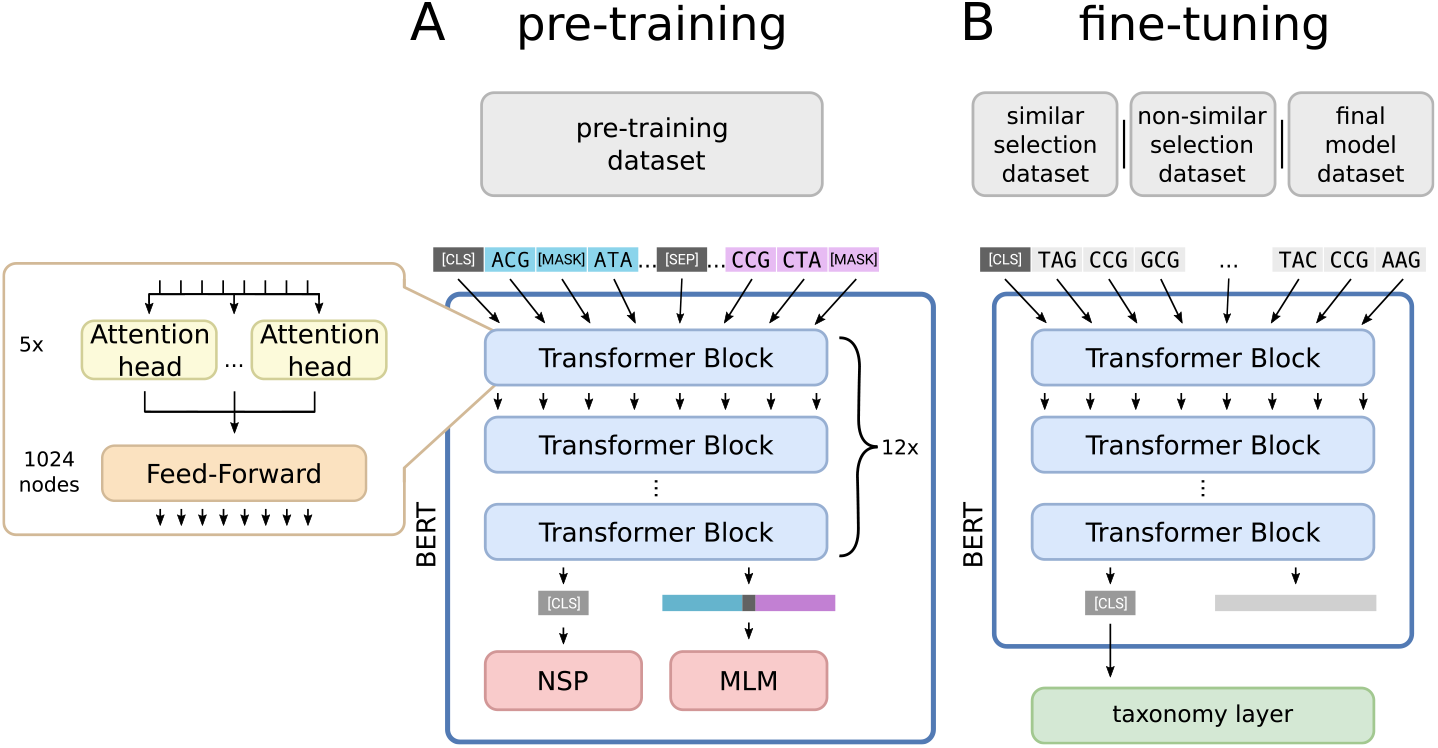
The training process of the `BERTax` model. **A** During pre-training, two separate sentences are used as input. Twelve stacked transformer blocks process this input, each consisting of five attention heads that determine which parts of the inputs are relevant to each other and a feed-forward layer that combines the results. The input, consisting of 502 tokens with 250 embedding dimensions each, is passed through all transformer blocks. Last, the NSP layer uses the classifier token `[CLS]` to predict if the second sentence is an original successor sentence or a replaced sentence. The MLM layer predicts all masked tokens from the remaining tokens. The evaluation of MLM and NSP evaluates the pre-training of the transformer blocks, and thus the quality of the learned language. After pre-training the pre-trained transformer blocks are used for the fine-tuning step. **B** A single sentence is used as input during fine-tuning with one of the three datasets (similar dataset – training and test data are phylogenetically closely related sequences, non-similar dataset – unknown genera are used as test dataset, and final model dataset – an expanded similar dataset). The pre-trained transformation blocks process this input, and only the `[CLS]` token is delivered to the taxonomy layers as output (see Fig. 4).

The length of the input is 502 tokens, composed of 500 tokens from the DNA sequence and two architecture-specific tokens to keep the sentences apart (see Fig. 1). This input is passed through all transformer blocks (see Fig. 2). As typical for an encoder architecture like BERT, the dimensionality of the sequence passed between the blocks and layers stays the same. The output of the pre-training architecture is comprised of the NSP-layer (i.e., the prediction whether or not the second sentence is a random replacement) and the MLM-layer with one output per input position (502) and distinct token (69) for predicting the 3-mer of a masked position.

### 2.2 Data

Archaean and eukaryotic genomes were retrieved from NCBI using `ncbi-genome-download`^1^, viral and bacterial genomes were downloaded from NCBI manually. The list of genomes including the respective assembly versions is provided in the supplement Tab. S2.

#### Pre-training dataset

For each superkingdom, we extracted one million fragments of length 1 500 nt from the downloaded genomes. To obtain a taxonomically balanced dataset, we grouped the genomes by the taxonomic rank ‘order’ and extracted fragments evenly distributed from these groups. Low-quality fragments with more than 0.08 % ambiguous characters were discarded and replaced. To reduce redundancy, we clustered fragments by sequence identity using `MMseqs2-linclust` version `11.e1a1c` [22] with a threshold of 80 %, resulting in 939 357 eukaryotic, 764 161 bacterial, 535 153 archaean and 253 803 viral fragments (see Tab. 1, pre-training dataset). The varying degree of reduction across superkingdoms can be explained by (a) the different number of taxa in each superkingdom, (b) the number of genomes in databases, (c) and the genome sizes. We refer to this dataset as *pre-training dataset*, used for pre-training of `BERTax`. As pre-training is unsupervised, the target classes of the taxonomic classification objective are not used and model weights are trained solely on the generic tasks MLM and NSP.

**Table 1:** The development and the number of fragments/samples of the datasets. We extracted one million taxonomically balanced fragments of length 1 500 nt from the downloaded genomes for each superkingdom. The resulting fragments were clustered by 80 % sequence identity forming the **pre-training dataset** used as input for pre-training. Next, we selected from the pre-training dataset all phyla with at least 10 000 fragments. We partitioned these fragments into a test set (2 000 sequences per phylum), training set (95 % of non-test set), and validation set (5 % of non-test set). For this we used two different approaches, similar selection and non-similar selection (see Fig. 2, 3), resulting in two different datasets, **similar dataset** and **non-similar dataset**. Since the genomic diversity of eukaryotes (939 357) and bacteria (765 161) seem to be less covered compared to archaea (535 153) and viruses (253 803) (see pre-training dataset), we extracted another two million fragments and clustered again by sequence similarity. Next, we selected all superkingdoms, phyla, and genera with at least 10 000 fragments. However, all clades with less than 10 000 fragments are combined into a new class ‘unknown’ for each taxonomic rank. Therefore, the number of fragments does not decrease. All fragments were subsequently divided via similar selection into test set, training set, and validation set (see proportions above). This process results in the **final-model dataset**.

| superkingdom | eukaryotes | bacteria | archaea | viruses |
| --- | --- | --- | --- | --- |
| # extracted fragments | 1 000 000 | 1 000 000 | 1 000 000 | 1 000 000 |
| after clustering | 939 357 | 764 161 | 535 153 | 253 803 |
| resulting dataset | **pre-training dataset** |  |  |  |
| after phylum selection | 873 873 | 690 984 | 535 153 | 228 574 |
| after partitioning* |  |  |  |  |
| similar selection | 873 873 | 690 984 | 535 153 | 228 574 |
| non-similar selection | 854 524 | 684 813 | 532 214 | 227 265 |
| resulting dataset | **similar & non-similar dataset** |  |  |  |
| # extracted fragments | 2 995 950 | 3 000 009 | 1 000 000 | 1 000 000 |
| after clustering | 2 707 781 | 1 903 183 | 535 153 | 253 803 |
| after clade selection | 2 707 781 | 1 903 183 | 535 153 | 253 803 |
| after partitioning* | 2 707 781 | 1 903 183 | 535 153 | 253 803 |
| resulting dataset | **final-model dataset** |  |  |  |

\*into training, validation and test set

#### Preparing data for fine-tuning

For fine-tuning we sorted the fragments from the pre-training dataset into classes according to the phylum the fragment originated from. We selected only classes with at least 10 000 fragments. All fragments of the selected classes form the input data, which we refer to as ‘samples’ as common in machine learning.

This results in 30 phylum classes with 2 328 584 samples in total. We partition this data into training, validation and test set. The training set is used to train the neural network. The validation set is used to prevent over-fitting, by evaluating the neural network’s loss (the distance to the optimal solution) after each training epoch, which allows “early stopping”, i.e., terminating training when the loss of the validation set does not further decrease. The test set is used to evaluate the performance of the neural network after finishing the training.

#### Three evaluation datasets

We used three different evaluation datasets. The first and second datset are compared to show the power of `BERTax` when unknown sequences are part of the test set. This is achieved by using different strategies to partition our data into training, validation and test sets (see Fig. 3).

**Figure 3:**
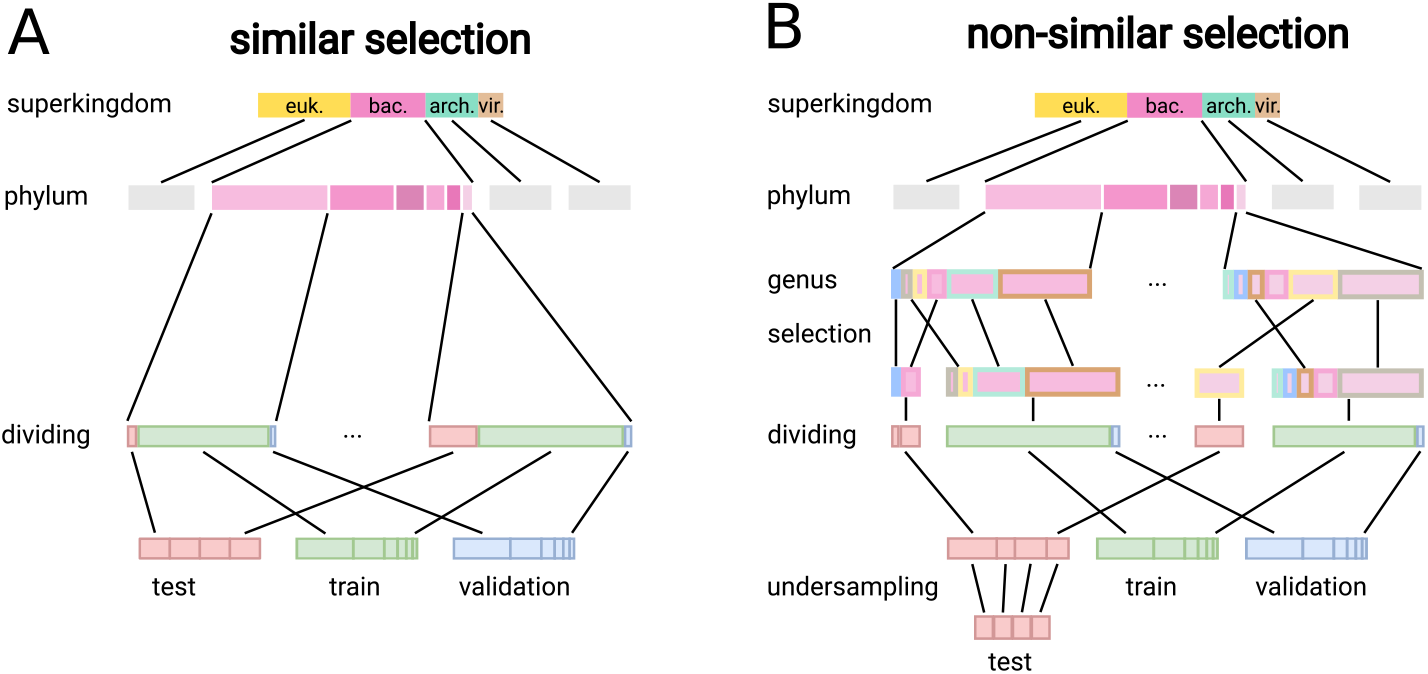
Data preparation of the test, training, and validation set via similar selection and non-similar selection. **A** The similar selection uses 2 000 randomly selected samples per phylum as a test set. All remaining samples of each phylum are divided into 95 % as training set and the remaining 5 % as validation set. **B** The non-similar selection selects per phylum one or more entire genera as test set (in comparison to the similar selection, where no restriction in this regard is made). This subset is the combination of genera with the number of samples closest to 2 000. The test set is balanced by undersampling, reducing the size per phylum to the size of the smallest subset. As for the similar selection all remaining samples of each phylum are divided into 95 % as training set and the remaining 5 % as validation set.

The comparison of the first and third dataset displays an increase of accuracy for all classes, although only classes are enriched, that are underrepresented considering how many data are available.

The first evaluation dataset is called **similar dataset**. As test set, we randomly select 2 000 samples per phylum. From the remaining samples we select 95 % of the samples of each phylum as training set and the remaining 5 % as validation set (see Fig. 3A).

The second evaluation dataset is called **non-similar dataset**. The goal of non-similar selection is to simulate sequences without (taxonomically) closely related sequences in the target database, simulating ‘unknown’ sequences. For this purpose, a genus separation between training and test set is employed, i.e., for no sample in the test set exists a sample of the same genus in the training set (see Fig. 3B). To achieve this, we split each phylum into its genera. As test set, we select a subset of these genera with about 2 000 samples; more precisely, the subset with the number of samples closest to 2 000. The test set is balanced by undersampling randomly: we keep 1 780 samples per phylum (according to the smallest subset). From the remaining samples in each phylum, we again select 95 % for the training set and the remaining 5 % as validation set.

The third evaluation dataset is referred to as the **final-model dataset**. The initial redundancy of the extracted eukaryotic and bacterial fragments is very low, indicated by the low reduction of fragments after clustering the pre-training dataset (see Tab.1). Therefore, the genomic diversity of the two superkingdoms is probably underrepresented. Thus, for fine-tuning our final model which is used in the downloadable version of `BERTax`, we extracted an additional two million fragments for those two superkingdoms and clustered again by sequence similarity. With this dataset, a more complete snapshot of the genomes is provided, with a wider textual and taxonomic diversity.

For this dataset we sorted the fragments into classes according to the superkingdom, phylum and genus the fragment originated from. Again, we selected only classes with at least 10 000 fragments. Remaining classes are moved to an additional class ‘unknown’, which is introduced for each taxonomic rank (i.e. ‘unknown superkingdom’, ‘unknown phylum’, ‘unknown genus’). With this, the number of classes per taxonomic rank is five for the rank superkingdom, 44 for phylum and 156 for genera. We partition this dataset into training, validation and test set using similar selection as described above (see Fig. 3A).

### 2.3 Fine-tuning: taxonomic classification

Fine-tuning is used for the training of the pre-trained model on the problem of interest. In our case this is the prediction of phylogenetic taxa, by classification of sequences into different classes, representing different taxa. While fine-tuning, each sample is converted to its full length of 1 500 nucleotides into the corresponding 500 tokens and the classification token `[CLS]` (representing the whole sequence). Because the required sequence length is 502, one additional padding token (`[PAD]`) has to be added to the end. The pre-trained transformer blocks process these 502 tokens, and from the resulting output, the `[CLS]` token is used by the taxonomy layer/s to predict the taxonomy (see Fig. 2B and Fig. 4). Next, the weights and biases of all layers, including the pre-trained transformer blocks, are adapted for a more precise prediction given this specific input.

**Figure 4:**
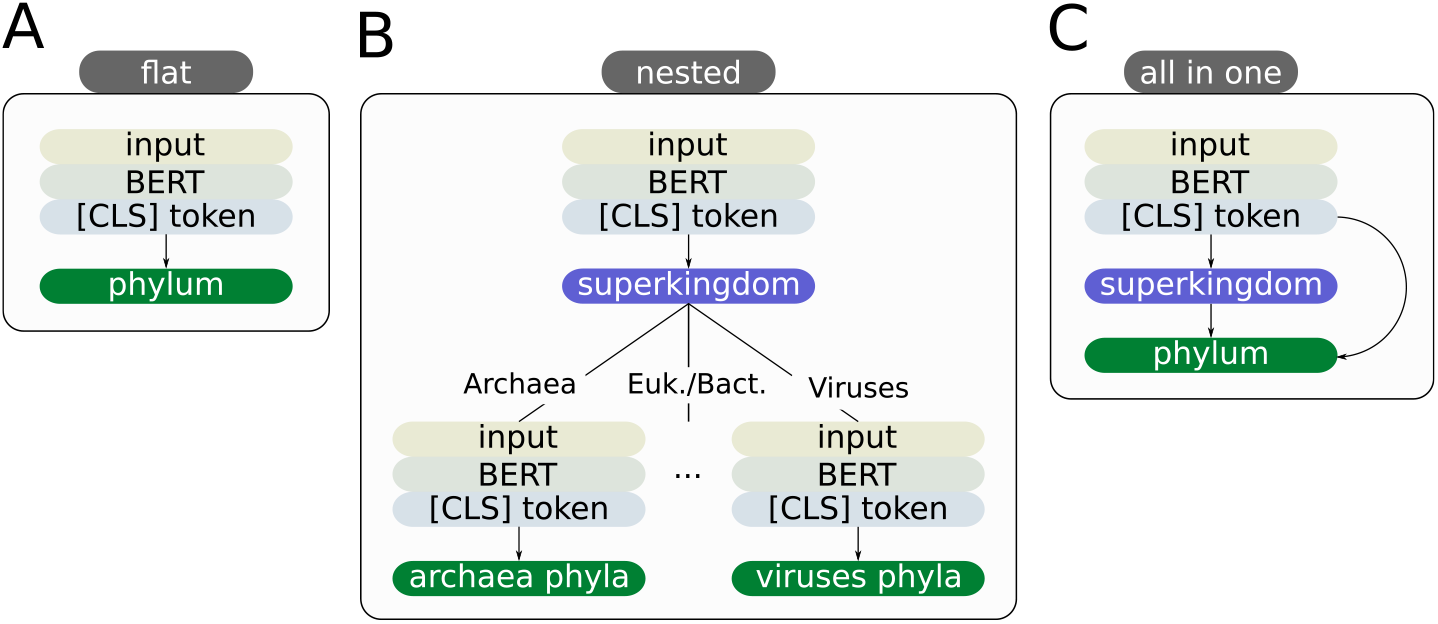
The evaluated architectures. Note this comparison was made on the taxonomic ranks, superkingdom, and phylum. However, the final `BERTax` model can furthermore predict the genus. (A) The **flat architecture** consists of a BERT model which is trained to directly predict the lowest taxonomic rank (the phylum), from which higher taxonomic ranks are inferred. (B) The **nested architecture** consists of multiple BERT models that are nested such that the top model takes each input and classifies the superkingdom and the sample is then further classified by the model that was trained to predict the phyla for that superkingdom. (C) The **all-in-one architecture** consists of a single BERT model which predicts all taxonomic ranks simultaneously. For this, all output layers (all taxonomic ranks) of the model have access to the BERT model itself and the output layer (the prediction) for higher taxonomic ranks.

We fine-tuned all models for a maximum of 16 epochs, employing early stopping (see supplement Tab. S3 for the exact number of epochs trained).

To avoid bias towards predicting the most frequent classes we balance the classes for fine-tuning by class weights, calculated for each taxonomic rank:

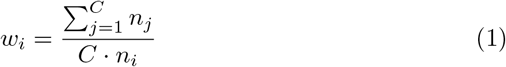

Here, *w_i_* is the class weight and *n_i_* the number of samples for class *i* = 1*, …, C*, applied to each sample. Using these weights allows preserving the complete amount of samples while ensuring that the influence of each class on the finetuning is balanced.

### 2.4 Architectures

We tested three different architectures based on the pre-trained BERT model, which differ in the adaptations of the output layers (see Fig. 4). The pre-trained model is used to obtain a learned high-dimensional embedding of the input sequence. Specifically, the classification token `[CLS]`, which is a single vector designed to contain the information of the whole input sequence, is the input to the new output layer. Each output layer uses the activation function `softmax`, which results in a probability distribution for each taxonomic rank, such that each prediction is associated with a confidence.

The **flat architecture** (see Fig. 4 A) is trained to directly predict the lowest taxonomic rank, which is the phylum in the comparison of the architectures. The flat architecture uses a simple dense layer with a node for each phylum.

The **nested architecture** (see Fig. 4 B) consists of multiple independent BERT models arranged in a tree-like manner. Each of these models is trained only on samples of its taxonomic group and uses a dense layer containing the same number of nodes as its subclasses. First, the root model predicts the superkingdom. Then, the sample is passed to the model of the subclass with the highest probability. This is continued until the lowest taxonomic rank is reached.

The **all-in-one architecture** (see Fig. 4 C) is a single BERT model which predicts all taxonomic ranks simultaneously. It uses the idea of Rojas-Carulla et al. [23] to provide the prediction of higher taxonomic ranks to lower prediction layers. For this, all output layers (taxonomic ranks) of the model have access to the BERT model itself and the output layers of all higher taxonomic ranks.

#### 2.4.1 Downloadable BERTax version

The downloadable `BERTax` version is built on the all-in-one architecture and trained on the *final-model dataset*. The downloadable tool additionally predicts the taxonomic rank genus.

`BERTax` was implemented in Python 3.7 and uses the Python packages scipy (1.6.1) [24], keras (2.4.3), tensorflow (2.4.1) [25], numpy(1.19.2) [26] and keras-bert (0.86.0). The visualization feature is based on bertviz (1.0.0). The source code as well as a conda package and docker container are available at https://github.com/f-kretschmer/bertax.

### 2.4.2 Comparison to other tools

`BERTax` is compared against the state-of-the-art database taxonomic classification approaches `Kraken2` [11], `sourmash` [12], `MMseqs2` [10], `Kaiju` [8], and `minimap2` [13]. Hereby `Kraken2` and `sourmash` use *k* -mers and minimizers for comparing the query to the reference database. `MMseqs2` uses *k* -mers and local alignments. `Kaiju` translates DNA input with the six possible reading frames into aminoacids and searches with Burrow-Wheeler transformation for similar entries in the reference database. `Minimap2` uses minimizers for the identification of seeds, which are further extended in a local alignment. For all tools we used the default parameters. However, we use a modified version of `MMseqs2 taxonomy`. More precisely, we are using `MMseqs2` with the parameters of `MMseqs2 taxonomy` (`--e-profile 0.001, -e 1`). Doing so, we get an E-value and thus a significance value that can be used to calculate the precision-recall curve and average precision, rather than assigning the same confidence to all predictions. With this approach we receive the same hits as for `MMseqs2` `taxonomy` for 99.94 % of all samples. We call this approach `MMseqs2 taxonomy`^*^. As state-of-the-art machine learning approach we tested `DeepMicrobes` [15]. In our evaluation we used the architecture and hyper-parameters evaluated as best by Liang *et al.* The architecture comprises LSTM-layers with self-attention that use *k* -mer embeddings (k=12) as input. The `DeepMicrobes` models are trained on exactly the same data as `BERTax`.

The deep-learning based method `GeNet` [23], developed for classifying bacteria, unfortunately could not be compared against, as it relies on downloading and subsequently binarizing training data on its own, which is highly impractical for our much bigger datasets. For `CNN-RAI` [27], only the source code of the tool is provided. Therefore, it is impossible to train the method on new data with reasonable effort, which is necessary for comparability.

`Seq2Species` [28] and `MetagenomicDC` [29] restrict the input to 16S sequences, which severely limits the approaches’ applicability, these tools are therefore not included in our comparisons. However, most of these approaches use Convolutional Neural Networks (CNN), which use combinations of short letter sequences (3-12 nt) for classification. This is similar to the use of *k* -mers in database approaches.

## 3 Results and Discussion

### 3.1 Best architecture based on simultaneous classifications of taxonomic ranks

We compared three different architectures (flat, nested and all-in-one, see Fig. 4) for predicting taxonomic classes. We used two different evaluation datasets (similar and non-similar dataset) for which we determine the accuracy of each architecture for predicting superkingdom and phylum (see Tab. 2). To prevent biasing the metric due to unbalanced data, we calculate the accuracy for each superkingdom class and use the mean over all superkingdom classes.

**Table 2:** Comparing different BERTax architectures (flat, nested, all-in-one) for taxonomic classification. The similar selection test set contains samples from the same genera as the training set. The non-similar selection test set contains samples from different genera as the training set. Each sample has been classified by superkingdom and phylum. The all-in-one architecture has the highest accuracy in all cases.

| architecture | similar dataset |  | non-similar dataset |  |
| --- | --- | --- | --- | --- |
|  | superkingdom | phylum | superkingdom | phylum |
| flat | 94.64 | 83.16 | 87.66 | 55.77 |
| nested | 90.31 | 73.12 | 86.95 | 50.12 |
| all-in-one | 94.78 | 85.55 | 88.95 | 60.10 |
| random | 25.00 | 3.33 | 25.00 | 3.33 |

Interestingly, the all-in-one architecture has the highest accuracy independent of the dataset (similar or non-similar dataset) and the taxonomic rank (superkingdom or phylum).

The all-in-one architecture provides the predicted likelihood of all classes of higher taxonomic ranks to lower prediction layers. This is advantageous compared to the flat architecture, as higher taxonomic classes indicate which subset of lower taxonomic classes is likely to contain the correct prediction. However, unlike the nested architecture, the higher taxonomic class classification does not categorically exclude some lower taxonomic classes. This has several advantages. Misclassification of higher taxonomic ranks (e.g. superkingdom) does not inevitably prevent the correct prediction of lower ranks. Further, the training of lower taxonomic ranks can also adjust the weights and biases of the neural network. Thus, not only large general features can be identified but also features of subgroups (see supplement Figure Fig. S2). Due to the relationships of the species’ genomes, these subgroups are lower taxonomic ranks. As a result of better subgroup prediction the prediction of higher taxonomic ranks advances as well. For further analysis, we only use the all-in-one architecture.

### 3.2 Comparison to other classification approaches

The database taxonomic classification approaches compared against either directly predict the taxonomic origin of a query (`Kraken2`, `sourmash`, `Kaiju`) or find the most similar sequence in a reference database (`MMseqs2` and `minimap2`), which can also be considered a prediction of taxonomic origin.

The average precision (AveP) for each tool is visualized in Fig. 5, exact values are listed in Tab. 3. The AveP ranges for prediction of superkingdoms and phylum of the similar dataset from 25.00 % and 3.33 % (`Kaiju`) to 97.14 % and 87.18 % (`DeepMicrobes`), respectively.

**Figure 5:**
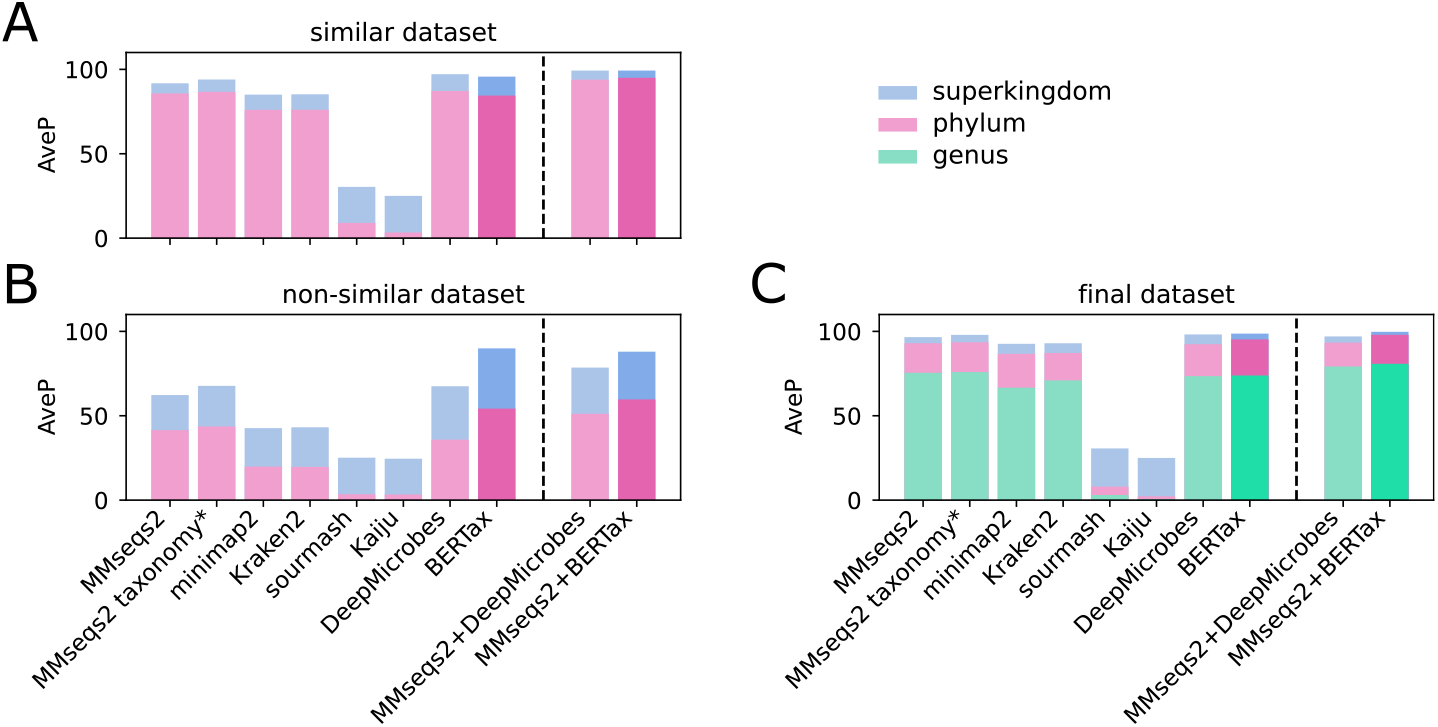
Visualization of the average precision (AveP) values on all three datasets. In the similar dataset, samples in the test set and the reference database (training set) can be from the same genus. In the non-similar dataset, samples in the test set do not have closely related (identical genus) samples in the reference database. The final dataset can contain samples in the test set and reference database of the same genus, like the similar dataset, but comprises of more samples. We queried by either superkingdom, phylum or genus (only final dataset). `MMseqs2 taxonomy`*: We used `MMseqs2` with the parameters of `MMseqs2 taxonomy` (`--e-profile 0.001, -e 1`)

**Table 3:**
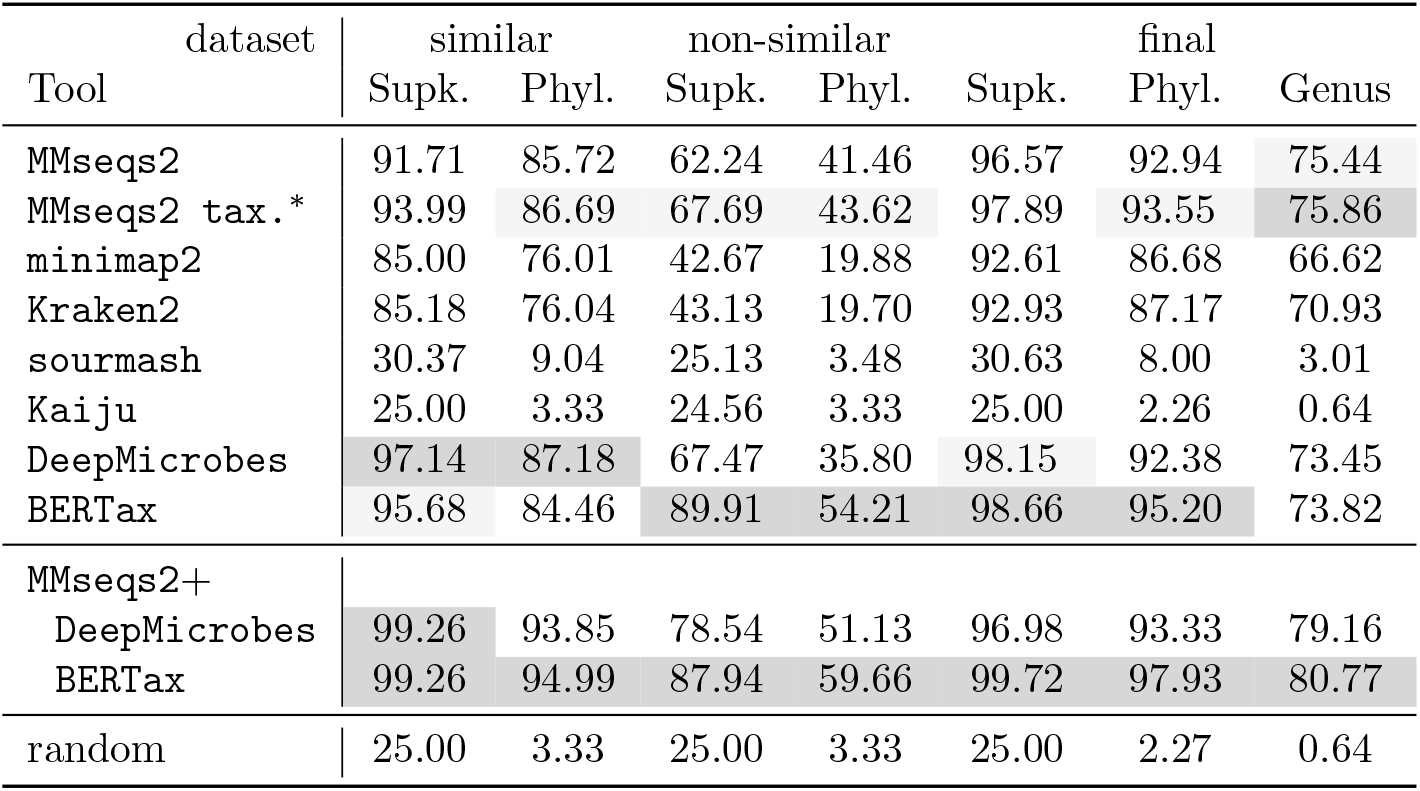
Comparison of the average precision (AveP) on all three datasets. For a description of the datasets, see e.g., Fig. 5 above. Taxonomic classification ranks considered are superkingdom (Supk.), phylum (Phyl.) or genus (only final selection dataset). The best performing tool is shown in dark grey and the second best in light grey.

Kaiju classified up to 100 % of queries as root or cellular organism, and thus higher than our highest taxonomic rank superkingdom. For these queries hits with diverse taxonomy were identified, for which the last common ancestor is ranked above the superkingdom. As Kaiju expects sequences to match annotated proteins from microbial reference genomes, the use of Kaiju as a general taxonomic classifier is not appropriate. However, our reference database contains a mixture of protein-coding and non-protein-coding sequences. Therefore, and because the reference database is bloated due to translation into six reading frames, false-positive hits are likely to occur. Resulting in the described hits with diverse taxonomy, and therefor the low AveP. The low performance is surprising, however, if the sequences used by a user match annotated proteins, we expect Kaiju to be highly effective.

Additionally, we devised a combination of classic database approaches with machine learning methods, which increases the overall recall while preserving the precision, resulting in more useful taxonomic information. We first predict the taxonomy by querying the reference database using `MMseqs2`, as it is a method with a high proportion of predicted samples and high accuracy (see supplement Tab. S1). We then predict the taxonomy of all unsuccessfully queried samples using `BERTax` and `DeepMicrobes` respectively. This strategy limits the amount of incorrectly predicted samples. Using this strategy `MMseqs2` + `BERTax` (99.26 % and 94.99 %) performs better than `MMseqs2` + `DeepMicrobes` (99.26 % and 93.85 %). This is to be expected as both `MMseqs2` and `DeepMicrobes` depend on *k* -mers for their prediction. Therefore, samples that are difficult to classify for `MMseqs2` tend to be also challenging to classify for `DeepMicrobes`, resulting in a limited higher precision. The combination of `BERTax` and `MMseqs2` outperforms all competing tools.

Real-world use cases always comprise a mix of sequences with and without similar representatives in the reference database. Therefore, it is reasonable to use a combined approach of a database approach and `BERTax`, as the classification performance is likely better, both recall- and precision-wise.

When we compare the classification performance of different tools, we typically observe a trade-off between accuracy and the proportion of samples classified (see supplement Tab. S1). If the classification tool uses a restrict threshold, we observe a very precise classification, however, a lower number of sequences could be classified. Whereas, when a high number of sequences is classified, usually it comes with the drawback of a less precise classification.

For further analysis see Fig. S3, which shows micro average Precision-Recall (PR) curves to visualize the trade-off between accuracy and proportion of classified sequences (see also supplement Tab. S1). PR curves reflect the quality of prediction independently of the sensitivity of the tested tool.

### 3.3 BERTax superior on *de novo* sequences

Unknown sequences are simulated by removing all sequences of one or several complete genera from the training set and corresponding sequences are only used in the test dataset (non-similar dataset). For this dataset `BERTax` outperforms with almost 90 % any other tool, ranging from 24.56 % to 67.69 % for superkingdom classification, see Tab. 3. Interestingly, `MMseqs2` + `BERTax` performs not as good as `BERTax` alone, hinting for a not stringent enough classification of `MMseqs2`. Instead of not classifying a sequence and passing it to the deep neural network, a misclassification leads to worse results, which could be changed by using more stringent standard parameters for `MMseqs2`. On phylum level, `BERTax` (with or without `MMseqs2`) outperforms all other tools albeit with a smaller margin than for superkingdoms. For both taxonomic ranks, `DeepMicrobes` performs worse than the best database approach, MMseqs2 taxonomy^*^.

These results are especially remarkable, since only a tiny fraction of organisms is so far described in databases and the power of `BERTax` outperforming any other classification tool for *de novo* sequences appears to be of utmost importance to the metagenome community.

### 3.4 Performance of final BERTax model

We evaluated the final `BERTax` model in comparison to other methods on the final-model dataset. This dataset differs from the similar dataset in that the number of eukaryotic and bacterial fragments is almost tripled. This results in an increased number of closely related sequences in the reference database. Expectedly, we observed higher average precision values for the final selection dataset compared to the similar dataset (see Tab. 3 and supplement Fig. S4). Comparing `BERTax` to the database approaches, it is performing better for superkingdom (99.72 %) and phylum (97.73 %) prediction. For the prediction of the lowest taxonomic rank (genus), `BERTax` achieves with 73.82 % comparable performance to the database approaches, however `MMseqs2` and `MMseqs2` `taxonomy`^*^ perform slightly better. Interestingly `BERTax` reached for all taxonomic ranks higher average precision than `DeepMicrobes`. This is in contrast to the comparable, yet smaller, similar dataset indicating an architectural advantage when predicting more taxonomic ranks while providing more training data.

The decrease in predictive ability for lower taxonomic ranks, also seen for the similar dataset and non-similar dataset, might be due to the lower number of available training examples per class and the chosen architecture. Further it is plausible that lower taxonomic ranks provide more advantages for the correct prediction of higher taxonomic ranks than *vise versa* (see supplement Fig. S2).

The combined approach of `MMseqs2` + `BERTax` is again the best performing one (see Table 3 and supplement Fig. S4). `MMseqs2` + `BERTax` reached on the final dataset an increase in aveP of 0.46 percentage points for the rank superkingdom and almost 3 percentage points for the rank phylum (see Tab. 3) compared to the similar dataset.

Although only Eukaryotes and Bacteria were increased in their training data, for all four superkingdoms the AveP of `BERTax` improved significantly, between 2.58 to 3.24 percentage points, see supplement Tab. S4.

### 3.5 Peeking into the Black-Box

Although (deep) neural networks are a powerful tool, their behavior is often hard or impossible to interpret, resembling a “black box”. However, as BERT at its core uses a (self-)attention mechanism, it is possible for us to gain insights into the feature relations and classification for given examples, see supplement Fig. S5 drawn with bertviz^2^[30]. Analyzing all inferred syntactic and semantic relations of DNA tokens presupposed by the model would be beyond the scope of this paper. However, some general remarks can be made regarding the interaction between attention heads in the same layer and in between whole layers: In general, weight patterns show a high degree of heterogeneity between attention heads, suggesting, that similar to natural languages, each attention head represents a different kind of syntactic relationship between the tokens[31]. The analysis of the visualization highlights the prominence of the special `[CLS]` token, whose encoding according to BERT’s design comprises the processed information of the whole sequence. A thorough analysis of the learned patterns and 3-mer relationships could be a promising starting point for further research.

## 4 Conclusion

Even today’s most comprehensive sequence databases miss significant portions of the total biodiversity. This incompleteness can result in large proportions of taxonomically unclassifiable DNA sequences. This is a common problem when using classical approaches, based on similarity to known sequences, on environmental samples. Here we present `BERTax` to tackle this task. `BERTax` is a tool for taxonomic classification of DNA sequences (reads, contigs, or scaffolds) using Deep Neural Networks. The ranks considered are superkingdom, phylum, and genus, although it seems natural to extend this to species and strain level in the future.

To the best of our knowledge, `BERTax` is the only taxonomic classification method not based on local similarities between the query and the target or limited to the use of *k*-mer frequencies, as even other deep-learning methods such as `DeepMicrobes` are. This provides a likely explanation for the superior generalization ability of `BERTax` as compared to other tools.

While our method has a higher average precision than comparable methods, a combination of a sensitive classic approach, like `MMseqs2`, and `BERTax` further improves the prediction performances, reaching unmatched sensitivity and specificity. The great power of `BERTax`, can be observed especially for sequences with no closely related species in the target database or training dataset.

Due to its high sensitivity and specificity, relatively independent of closely related species in the training data, `BERTax` is very suitable for use with metagenomic samples, especially on sequences that could not be classified with database methods. Thus, `BERTax` reduces the “microbial dark matter” and promises a significant benefit for metagenomics. In general, the applications of `BERTax` are manifold, as it can be applied in diagnostics to bypass otherwise lengthy cultivation processes, for example, in the diagnostics of fungi. Here, sequencing can be used together with `BERTax` to classify very rapidly directly at read level to identify potential infections. For the detection of endogenous viral elements (EVEs), the windowed mode of `BERTax` can be used to analyze target genomes. All windows classified as viruses can be considered as potential EVEs. In phylogenetics, it could pose as an alternative basis for distance determination of genomes. Here, all genomes are classified with `BERTax`, and for example, the pairwise Manhattan distances of the class predictions are used as a metric.

Alongside other publications [32, 19, 33, 34], our results yet again emphasize the power of Natural Language Processing methods in the field of biological sequences. Furthermore, a self-attention-based method like BERT grants a different perspective on the structure of DNA sequences, allowing – in theory – to investigate *how* the network comes to its classification “decision”, providing a promising starting point for further research on the structural organization of DNA sequences.

## Footnotes

1 https://github.com/kblin/ncbi-genome-download/, version 0.2.12

2 https://github.com/jessevig/bertviz

