## Supplement for "BERTax: taxonomic classification of DNA sequences with Deep Neural Networks"

### Supplementary Information

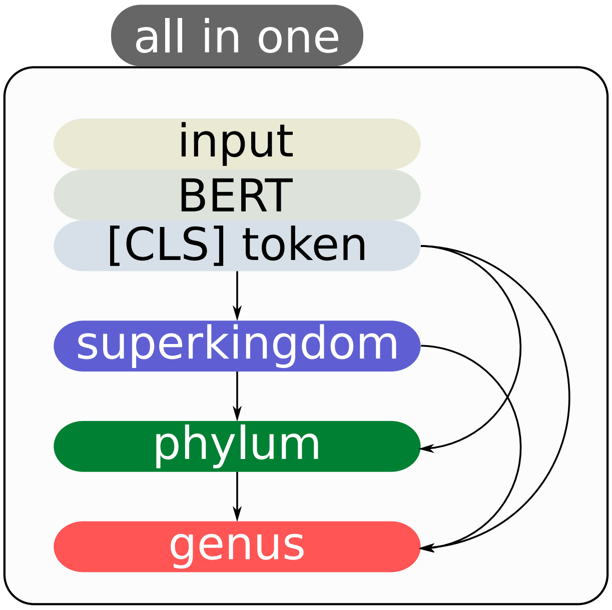


Figure S1: The final all-in-one architecture consists of a single BERT model which predicts all taxonomical ranks simultaneously (superkingdom, phylum, genus). For this all outputlayer (all taxonomical ranks) of the model, have access to the BERT model itself and the outputlayer (the prediction) for higher taxonomical ranks.

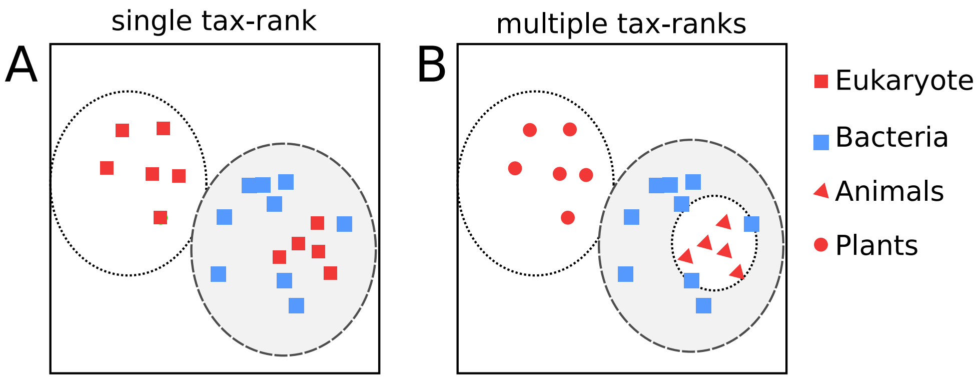


Figure S2: This example shows a possible distribution of training samples. The left figure (A) represents the training of a model on a single rank, here Superkingdom. All samples in the white circle are predicted as Eukaryota and in the gray circle as Bacteria. The figure shows that it can be difficult to correctly classify the red samples in the gray circle, as any adjustment to the model would initially decrease the prediction accuracy. The right figure (B) shows a model trained on Superkingdom and Kingdom simultaneously. Here, the training for Kingdom automatically adjusts the weights of the model to also address the Plants class, which ultimately also helps the Superkingdom prediction.

Table S1: Comparison of the accuracy (acc.) and the proportion (prop.) of the predicted samples per method on the similar and the non-similar dataset.

| Tool | similar dataset | | | | non-similar dataset | | | |
|---|---|---|---|---|---|---|---|---|
| | superkingdom | | phylum | | superkingdom | | phylum | |
| | acc. | prop. | acc. | prop. | acc. | prop. | acc. | prop. |
| MMseqs2 | 99.63 | 87.45 | 97.30 | 87.45 | 90.73 | 52.78 | 75.04 | 52.78 |
| MMseqs2_taxonomy* | 98.06 | 92.69 | 92.62 | 92.69 | 80.50 | 71.37 | 59.19 | 71.37 |
| minimap2 | 99.95 | 75.59 | 99.55 | 75.59 | 94.42 | 19.88 | 77.12 | 19.88 |
| Kraken2 | 99.82 | 76.11 | 98.82 | 75.95 | 90.71 | 21.56 | 68.26 | 21.08 |
| sourmash | 99.96 | 6.91 | 99.81 | 6.90 | 99.56 | 0.17 | 91.67 | 0.16 |
| Kaiju | 0 | 0 | 0 | 0 | 23.90 | 48.44 | 4.23 | 46.98 |
| DeepMicorbes | 96.68 | 100 | 87.72 | 100 | 68.39 | 100 | 41.95 | 100 |
| BERTax | 94.78 | 100 | 85.55 | 100 | 88.95 | 100 | 60.10 | 100 |
| MMseqs2 + DeepMicrobes | 99.10 | 100 | 94.03 | 100 | 78.20 | 100 | 56.28 | 100 |
| MMseqs2 + BERTax | 99.10 | 100 | 95.21.65 | 100 | 89.42 | 100 | 66.60 | 100 |
| random | 25.00 | 100 | 25.00 | 100 | 25.00 | 100 | 25.00 | 100 |

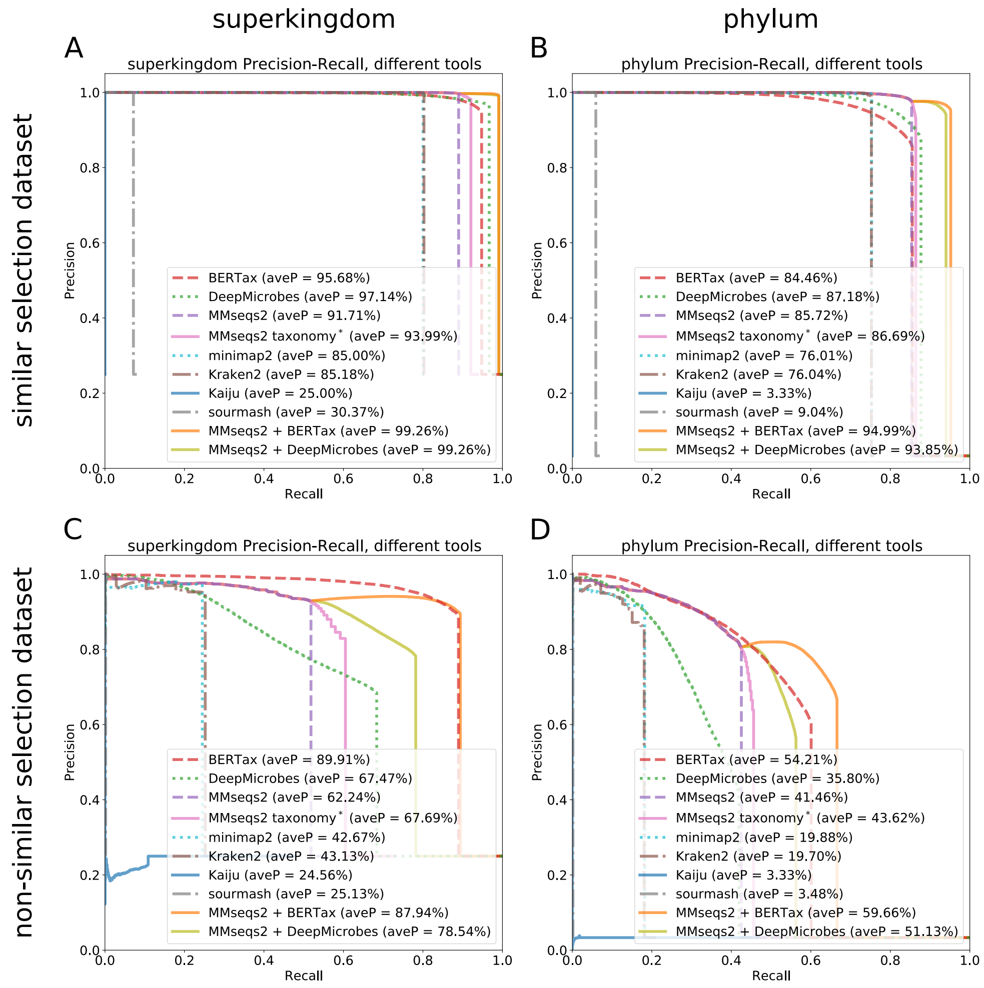


Figure S3: Precision-Recall curve on the similar and non-similar dataset, including samples with random guessing quality. The Recall (x-axes) indicates for how many samples the method could determine the correct superkingdom or phylum, at which precision (y-axes). If a method does not reach a recall of one, then not all samples were predicted from the method. Figures A and B show the results for the similar dataset, which comprises samples with the same genus in the query database (test set) as in the target database (training set). Figures C and D show the non-similar dataset, which comprises samples that do not have closely related (identical genus) samples in the query and target database.

### Fig. S3

For the similar dataset, differences in the performance at the rank superkingdom are only visible for high recall values (Fig. S3A). However, only the combination of `BERTax` and `MMseqs2` or `DeepMicrobes` and `MMseqs2` are able to achieve a recall of close to 1. For single approaches `BERTax` achieves the second highest average precision, after `DeepMicrobes`. At the level of phylum (Fig. S3B), `MMseqs2 + BERTax` is the best performing method achieving a near 1 recall. The precision of `BERTax` alone drops for lower recall values, although a high recall value is still achieved. In comparison, `MMseqs2` and `MMseqs2 taxonomy`* show a less sharp drop-off of precision at high recall values. Fig. S3C,D shows the superior performance of `BERTax` and `MMseqs2 + BERTax` on novel sequences. The difference in performance to the database approaches, and competing machine learning approaches is especially visible as theses approaches do not reach high recall values, both for classifying the rank of superkingdom and, to a lesser extent, phylum.

For all tools, we only consider the top prediction for each sample. This means that less likely predictions (e.g. the second most likely class) are not included. Moreover, not all tools return predictions for all samples. Thus, although the test set was balanced with regards to the taxonomic class, the set of predicted samples might not be. Therefore, we weight the micro average PR curves by the number of predicted samples for each taxonomic class. Note that for all samples in the test set `BERTax` provides taxonomic predictions.

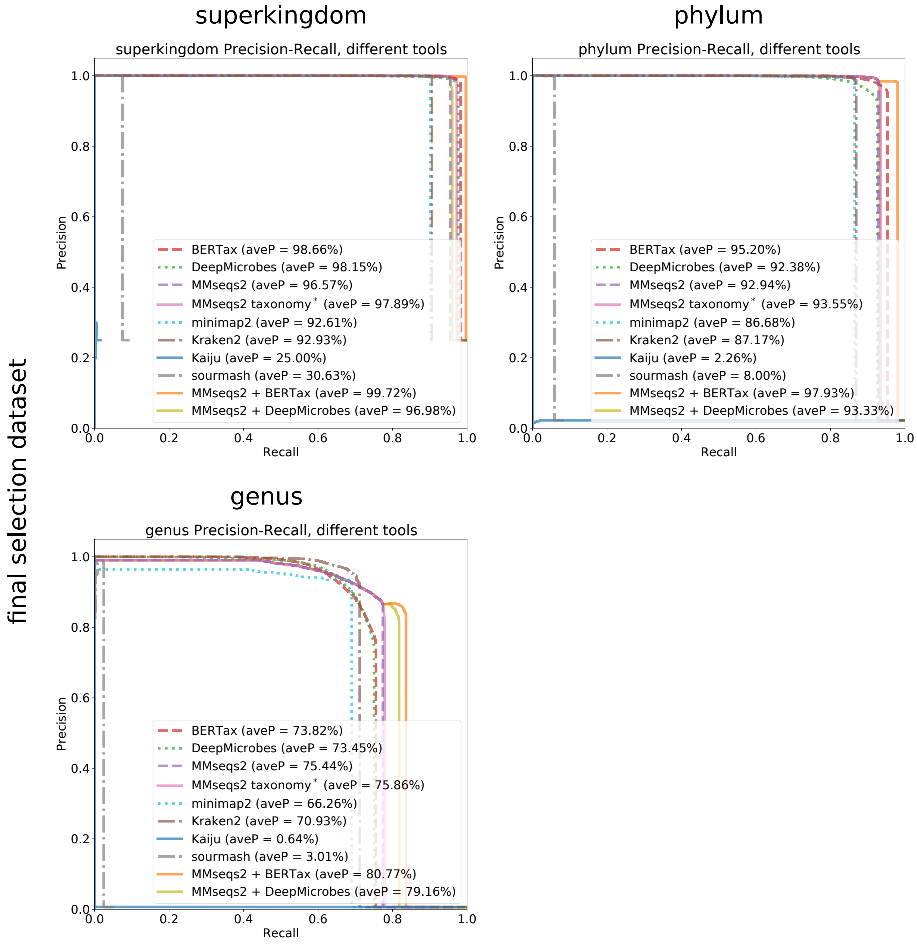


Figure S4: Precision Recall curves on the final selection dataset, including samples with random guessing quality. The Recall (x-axes) indicates for how many samples the method could determine the correct superkingdom or phylum, at which precision (y-axes). If a method does not reach a recall of one, then not all samples were predicted from the method.

### Fig. S4

The final dataset enables nearly all tools to achieve higher aveP compared to the similar dataset. This is expected as the dataset contains a higher number of similar sequences, which greatly enhances the chance of finding a similar sample for all database approaches and provides more training data for the machine learning approaches. At superkingdom and phylum level all approaches, except `Kaiju` and `sourmash` reach a very high aveP of over 85 %. However, `BERTax` and `MMseqs2 + BERTax` surpass all comparable approaches, reaching near perfect aveP results with 98.66 % and 99.72 % on superkingdom level and 95.20 % and 97.93 % of phylum level. On genus level `BERTax` is the third best approach after `MMseqs2` and `MMseqs2 taxonomy`*. However, if used in the combined fashion, `MMseqs2 + BERTax` outperforms all competing methods.

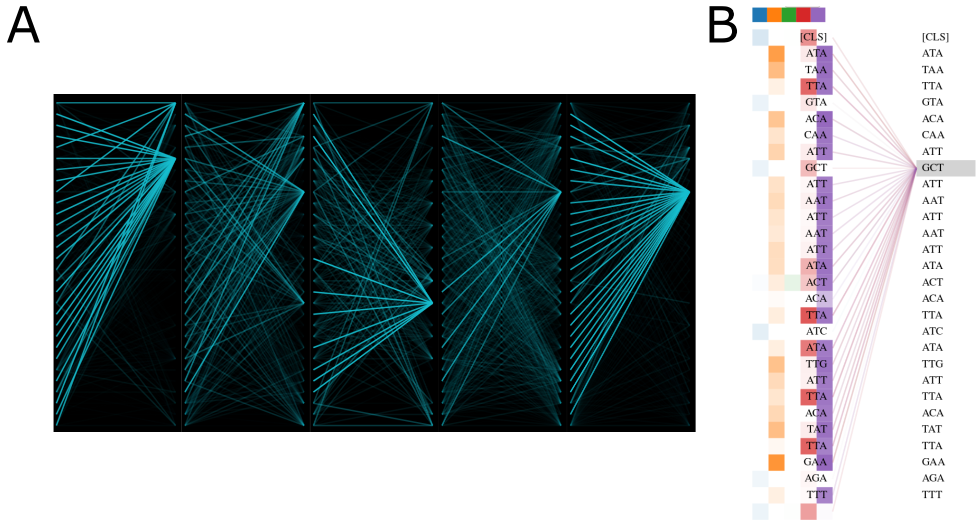


Figure S5: Visualization of the self-attention behaviour of `BERTax`. The sequence used here is a 90 nt fragment of the archaeum *Methanocorpusculum labreanum*, weights of the 10th layer are shown. **A:** Visualization of attention weights shows the coordinated distribution of different inter-token relationships across attention heads in the same layer. Attention heads are represented as graphs: Each of the five graphs represents one attention head in the 10th transformer layer. The attention heads represent independent relationships between the input tokens of the same sequence. The tokens are visualized from top to bottom as nodes, edges represent weights: thicker edges depict higher weights. **B:** An alternative visualization mode shows attention weights of the token 'GCT', the different attention heads are symbolized by differently colored rectangles.

Table S2: Genomes used to generate training and testing data with their respective assembly versions.

`genomes.xlsx`

Table S3: Number of epochs trained per evaluated `BERTax` architecture before early stopping, as the nested architecture comprises of multiple trained models the numbers can vary.

| architecture | flat | nested | all-in-one |
|---|---|---|---|
| epochs | 16 | 11-13 | 15 |

Table S4: Average precision (AveP) of the precision-recall curve for the final `BERTax` model compared to the model trained on the similar dataset.

| training dataset | Eukaryota | Bacteria | Archaea | Viruses |
|---|---|---|---|---|
| similar dataset | 96.07 | 96.41 | 95.40 | 94.73 |
| final dataset | 99.30 | 98.99 | 98.64 | 97.53 |
